# Growth factor receptor signaling inhibition prevents SARS-CoV-2 replication

**DOI:** 10.1101/2020.05.14.095661

**Authors:** Kevin Klann, Denisa Bojkova, Georg Tascher, Sandra Ciesek, Christian Münch, Jindrich Cinatl

**Author notes:** These authors contributed equally (Kevin Klann, Denisa Bojkova). These authors contributed equally (Christian Münch, Jindrich Cinatl). Corresponding author. (C.M.) and (J.C.).

## Abstract

SARS-CoV-2 infections are rapidly spreading around the globe. The rapid development of therapies is of major importance. However, our lack of understanding of the molecular processes and host cell signaling events underlying SARS-CoV-2 infection hinder therapy development. We employed a SARS-CoV-2 infection system in permissible human cells to study signaling changes by phospho-proteomics. We identified viral protein phosphorylation and defined phosphorylation-driven host cell signaling changes upon infection. Growth factor receptor (GFR) signaling and downstream pathways were activated. Drug-protein network analyses revealed GFR signaling as key pathway targetable by approved drugs. Inhibition of GFR downstream signaling by five compounds prevented SARS-CoV-2 replication in cells, assessed by cytopathic effect, viral dsRNA production, and viral RNA release into the supernatant. This study describes host cell signaling events upon SARS-CoV-2 infection and reveals GFR signaling as central pathway essential for SARS-CoV-2 replication. It provides with novel strategies for COVID-19 treatment.

## Introduction

Severe acute respiratory syndrome coronavirus 2 (SARS-CoV-2), a novel coronavirus, has been rapidly spreading around the globe since the beginning of 2020. In people, it causes coronavirus disease 2019 (COVID-19) often accompanied by severe respiratory syndrome (Chen et al., 2020). To conquer the global health crisis triggered by COVID-19, rapidly establishing drugs is required to dampen the disease course and relieve healthcare institutions. Thus, repurposing of already available and (ideally) approved drugs might be essential to rapidly treat COVID-19. Many studies for proposing repurposing of specific drugs have been conducted in the last months, but mostly remain computational without tests in infection models (Smith and Smith, 2020; Wang, 2020). In addition, they are hindered by the lack of knowledge about the molecular mechanisms of SARS-CoV-2 infection and the resulting host-cell responses required to allow viral replication. To rationally repurpose drugs, a molecular understanding of the infection and the changes within the host cell pathways is essential. Experimentally identifying viral targets in the cell allows candidate drugs to be selected with high confidence for further testing in the clinics to reduce the risks for patients resulting from tests with drugs lacking *in vitro* validation.

Growth factor receptor (GFR) signaling plays important roles in cancer pathogenesis and has also been reported to be crucial for infection with some viruses (Beerli et al., 2019; Kung et al., 2011; Zhu et al., 2009). GFR activation leads to the modulation of a wide range of cellular processes, including proliferation, adhesion, or differentiation (Yarden, 2001). Various viruses, such as Epstein-Barr virus, influenza, or hepatitis C, have been shown to use the epidermal growth factor receptor (EGFR) as an entry receptor (Eierhoff et al., 2010; Kung et al., 2011; Lupberger et al., 2011). In addition, EGFR activation can suppress interferon signaling and thus the antiviral response elicited in respiratory virus diseases, for instance influenza A and rhinovirus (Ueki et al., 2013). Activation of GFR signaling might play an important role also in other respiratory viruses, such as SARS-CoV-2.

In the last years, it has been shown for many viruses that modulation of host cell signaling is crucial for viral replication and might exhibit strong therapeutic potential (Beerli et al., 2019; Pleschka et al., 2001). However, how SARS-CoV-2 infection changes host cell signaling has remained unclear. We recently established an *in vitro* cell culture model of SARS-CoV-2 infection using the colon epithelial cell line Caco-2, which is highly permissive for the virus and commonly used for the study of coronaviruses (Herzog et al., 2008; Ren et al., 2006). Here, we determine changes in the cellular phospho-protein networks upon infection with SARS-CoV-2 to gain insight into infection-induced signaling events. We found extensive rearrangements of cellular signaling pathways, particularly of GFR signaling. Strikingly, inhibiting GFR signaling using prominent (anti-cancer) drugs – namely pictilisib, omipalisib, RO5126766, lonafarnib, and sorafenib – prevented SARS-CoV-2 replication *in vitro*, assessed by cytopathic effect and viral RNA replication and release. These compounds prevented replication at clinically achievable concentrations. Due to their clinical availability, these drugs could be rapidly transitioned towards clinical trials to test their feasibility as COVID-19 treatment option.

## Results

### Phospho-proteomics of cells infected with SARS-CoV-2

To evaluate changes in intracellular signaling networks brought about by SARS-CoV-2 infection, we quantified phospho-proteome changes 24 hours after infection (Figure 1A). Caco-2 cells were mock-infected or infected with SARS-CoV-2 patient isolates (in biological quintruplicates) for one hour, washed, and incubated for 24 hours before cell harvest. Extracted proteins were digested and split to 1) carry out whole-cell proteomics of a tandem mass tag (TMT) 10-plex samples using liquid chromatography synchronous precursor selection mass spectromety (LC-SPS-MS^3^), or 2) use Fe-NTA phosphopeptide enrichment (achieving 98% enrichment) for phospho proteome analyses of a TMT 10-plex analyzed by LC-MS^2^. We identified and quantified 7,150 proteins and 15,392 different phosphopeptides for a total of 15,040 different modification sites (Figure 1B, C, S1, and Table S1, S2). The main fraction of phosphopeptides were modified serines (86.4%), followed by threonine (13.4%), and tyrosine (0.2%) (Figure 1D). Upon infection, 2,197 and 799 phosphopeptides significantly increased or decreased respectively (log_2_ FC > 1, p value < 0.05).

**Fig. 1.**
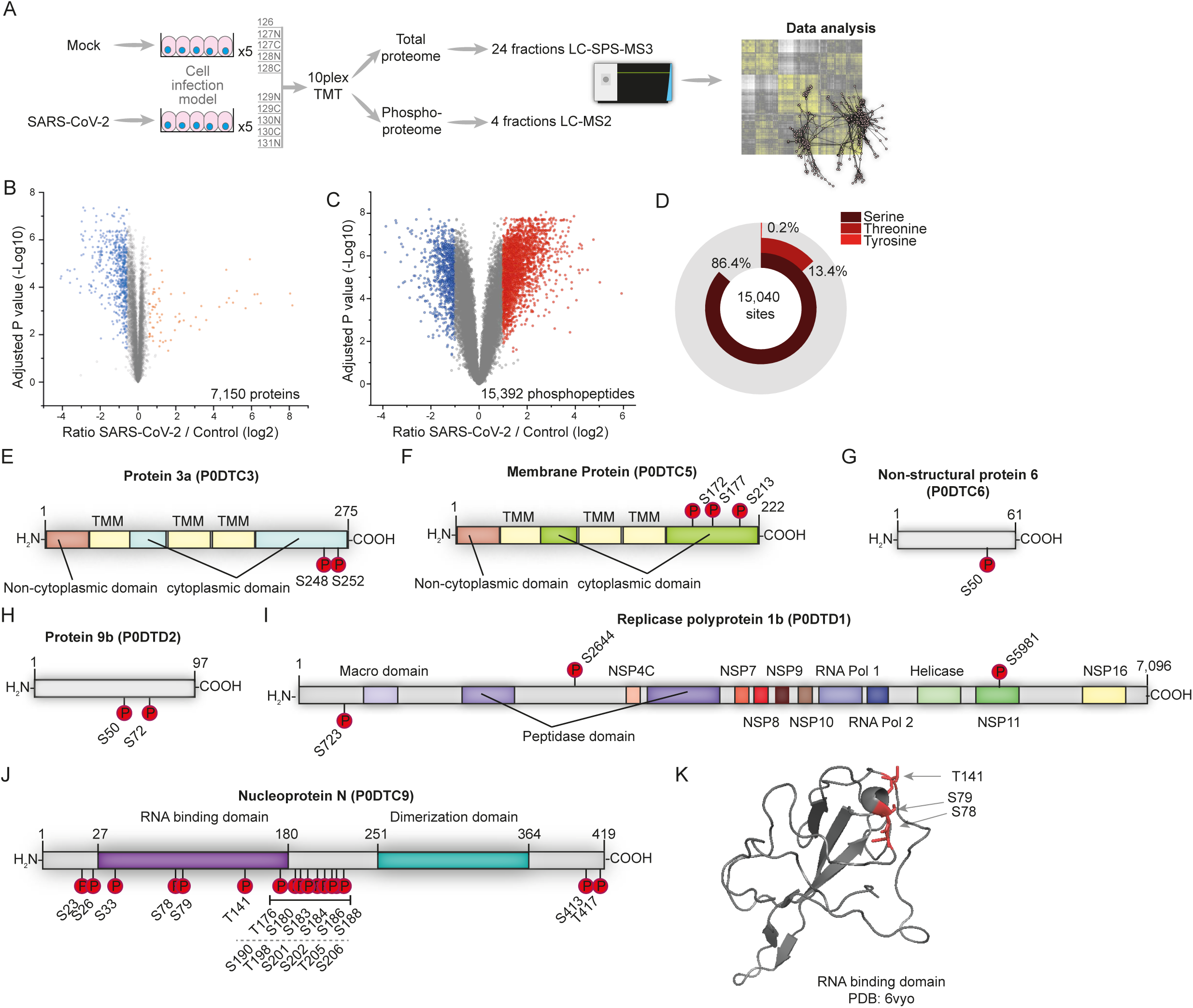
Phospho proteomic profiling of SARS-CoV-2 infected cells. (**A**) Experimental scheme. Caco-2 cells were infected with SARS-CoV-2 for one hour, washed and incubated for additional 24 hours. Proteins were extracted and prepared for bottom-up proteomics. All ten conditions were multiplexed using TMT10 reagents. 250 μg of pooled samples were used for whole cell proteomics (24 Fractions) and the remainder (~1 mg) enriched for phosphopeptides by Fe-NTA. Phosphopeptides were fractionated into 8 fractions and concatenated into 4 fractions. All samples were measured on an Orbitrap Fusion Lumos. (**B**) Volcano plot showing fold changes of infected versus mock cells for all 15,392 quantified phosphopeptides. P values were calculated using an unpaired, two-sided student’s t-test with equal variance assumed and adjusted using the Benjamini Hochberg FDR method (N = 5). Orange or blue points indicate significantly increased or decreased phosphopeptides, respectively. (**C**) Volcano plot showing differences between SARS-CoV-2 and mock infected cells in total protein levels for all 7,150 quantified proteins. P values were calculated using an unpaired, two-sided student’s t-test with equal variance assumed and adjusted using the Benjamini Hochberg FDR method (N = 5). Orange or blue points indicate significantly increased or decreased phosphopeptides, respectively. (**D**) Distribution of phosphorylation sites identified across modified amino acids. See also Figure S1 and Tables S1 and S2. (**E – K**) Domain structures of SARS-CoV-2 proteins predicted by InterPro. Identified phosphorylation sites are indicated.. Protein 3a (**E**), Membrane Protein M (**F**), Non-structural protein 6 (**G**), Protein 9b (**H**), Replicase Polyprotein 1b (**I**) and Nucleoprotein N (**J**). (**K**) X-ray structure of the RNA binding domain (PDB: 6vyo, residues 47-173) with identified phosphorylation sites marked in red.

Viral proteins are produced in the host cell and underlie (and often require) post-translational modification (PTM) by host cell enzymes (Wu et al., 2009). Accordingly, we assessed viral proteins phosphorylated in the host cell. We identified 33 modification sites on 6 different viral proteins (Figure 1E-J). Possible functions of the observed modifications largely remain unclear due to a lack of understanding of their molecular function and regulation. SARS-CoV-2 protein 3a was phosphorylated on the luminal side of this transmembrane protein (Figure 1E) Membrane protein M was phosphorylated at three serines in close proximity, at the C-terminal, cytoplasmic region of the protein (Figure 1F), suggesting a high-activity modification surface. SARS-CoV-1 protein 6 was described to accelerates infections in murine systems (Tangudu et al., 2007). We found a single phosphorylation of the SARS-CoV-2 protein homologue non-structural protein 6 in host cells (Figure 1G) Protein 9b was modified at two sites (Figure 1H). However, its function in SARS-CoV-1 or SARS-CoV-2 remains unknown. Polyprotein 1b is a large 7,096 amino acid protein heavily processed to generate distinct proteins in SARS-CoV-1 (Tangudu et al., 2007). We found polyprotein 1b to be modified at three residues, two in a region of unknown function and one in the non-structural protein 11 (NSP11) part of the protein (Figure 1I). Our data cannot distinguish whether phosphorylation occurred before or after cleavage and whether phosphorylation may affect processing. SARS-CoV-2 nucleoprotein was heavily phosphorylated (Figure 1J). Mapping phosphosites to the structure (residues 47 to 173, PDB: 6vyo) revealed a small surface region, suggesting specific regulation and interaction changes (Figure 1K). To reveal host kinases potentially phosphorylating viral proteins, we bioinformatically assessed identified phosphorylation motifs using NetPhos 3.1 and GPS5(Blom et al., 2004; Wang et al., 2020) (Table S3). Some motifs present in nucleoprotein were predicted to be modified by CMGC kinases. Among several others, casein kinase II (CK2) kinases are part of the CMGC family and have been independently identified as interaction partners of the nucleoprotein, when expressed in cells (Gordon et al., 2020). Inhibition of CK2 kinases, could be employed to study possible functional interactions between kinase and viral protein.

Taken together, we identified extensive changes in phosphorylation of host and viral proteins after SARS-CoV-2 infection. The role of viral protein modifications remain unclear. However, targeting the corresponding host kinases may offer new treatment strategies.

### Signaling pathways modulated upon infection

To identify the key host signaling pathway networks modulated by infection, we carried out protein-protein co-regulation analysis on all proteins quantified in phosphorylation and total protein level. We first standardized phosphorylation and total protein levels by individual Z-scoring to compare the different datasets. Subsequently, to merge phosphorylation and proteome data, we collapsed all phospho-sites for each protein into one average profile and calculated combined Z-scores. Patterns of co-regulation were identified using protein-protein correlation and hierarchical clustering (Figure 2A). The dynamic landscape of the proteome revealed three main clusters of co-regulated proteins, each one representing different sets of pathways (discussed in detail below):

**Fig. 2.**
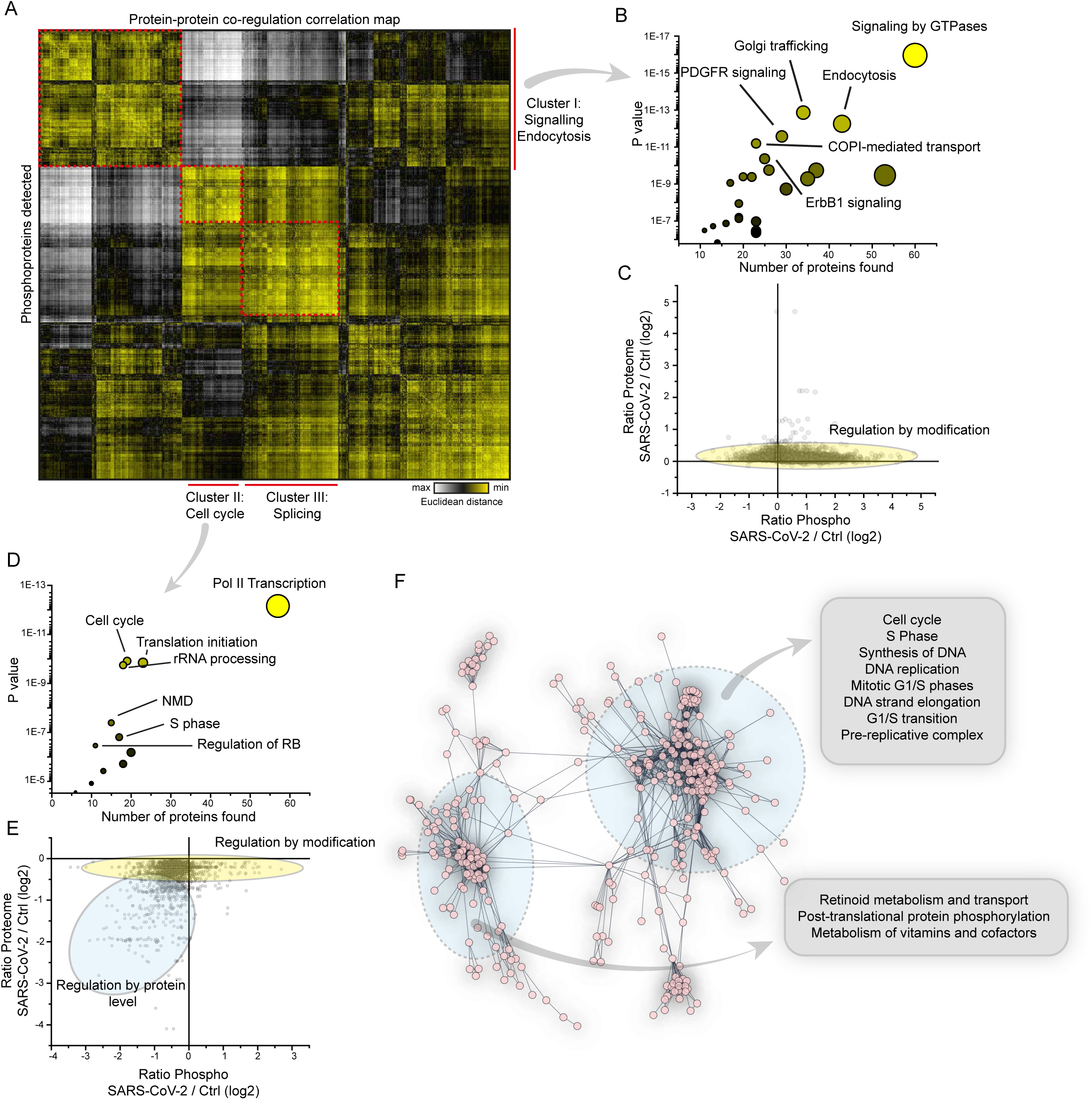
Correlation of co-regulated proteins identifies cellular signaling pathways modulated upon infection. (**A**) Correlation map of all detected phospho-proteins indicating Euclidean distance between proteins. To determine correlation, Z-scores of phospho-peptides and total protein levels were added and all peptide values for one protein collapsed into an average Z score. Correlation clustering was performed by Euclidean distance on combined Z scores for all conditions. Red dashed line indicates main clusters found and identified. (**B**) Reactome pathway enrichment of proteins found in Cluster I in (**A**). Shown are the number of proteins identified in the respective cluster versus statistical significance of enrichment. Circles are increasingly sized according to the number of proteins found in the pathway. (**C**) Scatter plot showing fold changes of phospho-peptides compared to fold changes of total protein levels. The yellow oval indicates peptides for which phosphorylation is not driven by changes in protein abundance. (**D**) Reactome pathways found enriched in Cluster II in (**A**). Analyses and presentation as in (**B**). (**E**) Scatter plot showing correlation between fold changes of phosphopeptides compared to fold changes of total proteins levels. Two subsets of phosphopeptides were detected: one was mainly regulated by differential modification (indicated in yellow), the second by changes in protein abundance. (**F**) STRING network analysis of proteins decreased in total protein levels (Figure 1C). Inserts indicate pathways found in the network

The first cluster was mainly comprised of receptor signaling and endocytic pathways (Figure 2B). Prominent among these pathways were platelet derived growth factor receptor (PDGFR), ErbB1 (EGFR) signaling, metabolism, and various pathways associated with vesicle trafficking (Table S4). As changes in phospho-peptide abundance can represent different ratios in phosphorylated versus non-phosphorylated peptide or a change in protein abundance (with the same ratio of protein being phosphorylated), we integrated our phospho-proteome dataset with total proteome data (Figure 2C). In contrast to the extensive changes observed in the phospho-proteome, no general changes were observed for the total proteome (Figure 2C, Table S2). Thus, phosphorylation changes were induced by signaling activity alteration resulting in increased phosphorylation and not due to protein abundance differences.

The second cluster was mainly comprised of proteins decreased in phosphorylation and was highly connected to cell cycle and translation initiation (Figure 2D and Table S5). We reported recently that inhibition of cellular translation prevented SARS-CoV-2 replication in cells (Bojkova et al., 2020), consistent with regulation of translation by altering phosphorylation patterns. To further distinguish the regulations within this cluster, we correlated protein levels with differential phosphorylation abundance (Figure 2E) and found two groups of proteins: The first was contained translation related pathways (identified in Figure 2E) and was predominantly regulated by decreased modification. The second set of proteins was decreased in phosphorylation and on total protein level. The majority of proteins found in the second cluster belonged to diverse cell cycle pathways. Consistent with these findings, cell cycle pathways were also enriched in the set of proteins significantly decreased on protein level (Figure 2F, S2, and Table S6). Translation pathways were not regulated on protein level to this extent.

Analysis of the third cluster revealed signaling events of the splicing machinery (Table S7) possibly explaining previously observed changes in splicing machinery abundance upon SARS-CoV-2 infection (Bojkova et al., 2020). Consistent with previous literature (Grimmler et al., 2005; Ilan et al., 2017; Mathew et al., 2008; Mermoud et al., 1994), we therefore hypothesized that the host splicing machinery is extensively reshaped during viral infection. This finding further supports splicing as a potential therapeutic target, in agreement with decreased SARS-CoV-2 pathogenic effects when inhibiting splicing by pladeinolide B. Additionally, we found carbon metabolism among the pathways showing significantly increased phosphorylation upon SARS-CoV-2 infection (Table S4) in addition to previously described changes of total protein levels of enzymes part of glycolysis and carbon metabolism (Bojkova et al., 2020)(Figure S3).

Taken together, we showed that, during SARS-CoV-2 infection, specific rearrangements of signaling pathways were elicited in the cellular proteome. Regulation was mainly comprised of cellular signaling and translational pathways as well as proteins regulated not only by phosphorylation, but also in total protein abundance. Proteins exhibiting decreased protein levels were significantly enriched in cell cycle proteins.

### Drug-target network reveals growth factor signaling as potent therapy candidate

We observed over 2,000 phospho-peptides to be increased in abundance while their protein levels stay constant upon infection (Figure 2C and S4). This reveals differential modification activity (e.g. signaling events) for these phospho-proteins. For many kinases in cellular signaling pathways there are already approved drugs available. Hence, we investigated the potential to repurpose drugs to treat COVID-19 by mapping already available drugs via ReactomeFI to the set of proteins increased in phosphorylation. We filtered the network for drugs and direct targets and found EGFR as one of the central hits, including a number of regulated proteins in the downstream signaling pathway of EGFR (Figure 3A). These downstream targets are also regulated by other GFRs and could thus also be explained by their observed activation upon SARS-CoV-2 infection (Figure 2). 28 clinically approved drugs (largely used in cancer therapy) are already available to target EGFR or downstream targets. Indeed, we found a subnetwork of GFR signaling components remodeled (Figure 3B). We mapped identified members of GFR signaling and their respective phosphorylation differences upon SARS-CoV-2 infection (Figure 3C) revealing an extensive overall increase in phosphorylation of the whole pathway, including related components for cytoskeleton remodeling and receptor endocytosis. How GFR signaling might regulate SARS-CoV-2 infection is still matter for speculation. However, GFR signaling inhibition might provide a useful approach already implicated in SARS-CoV induced fibrosis therapy (Venkataraman and Frieman, 2017) and might be a viable strategy to treat COVID-19.

**Fig. 3.**
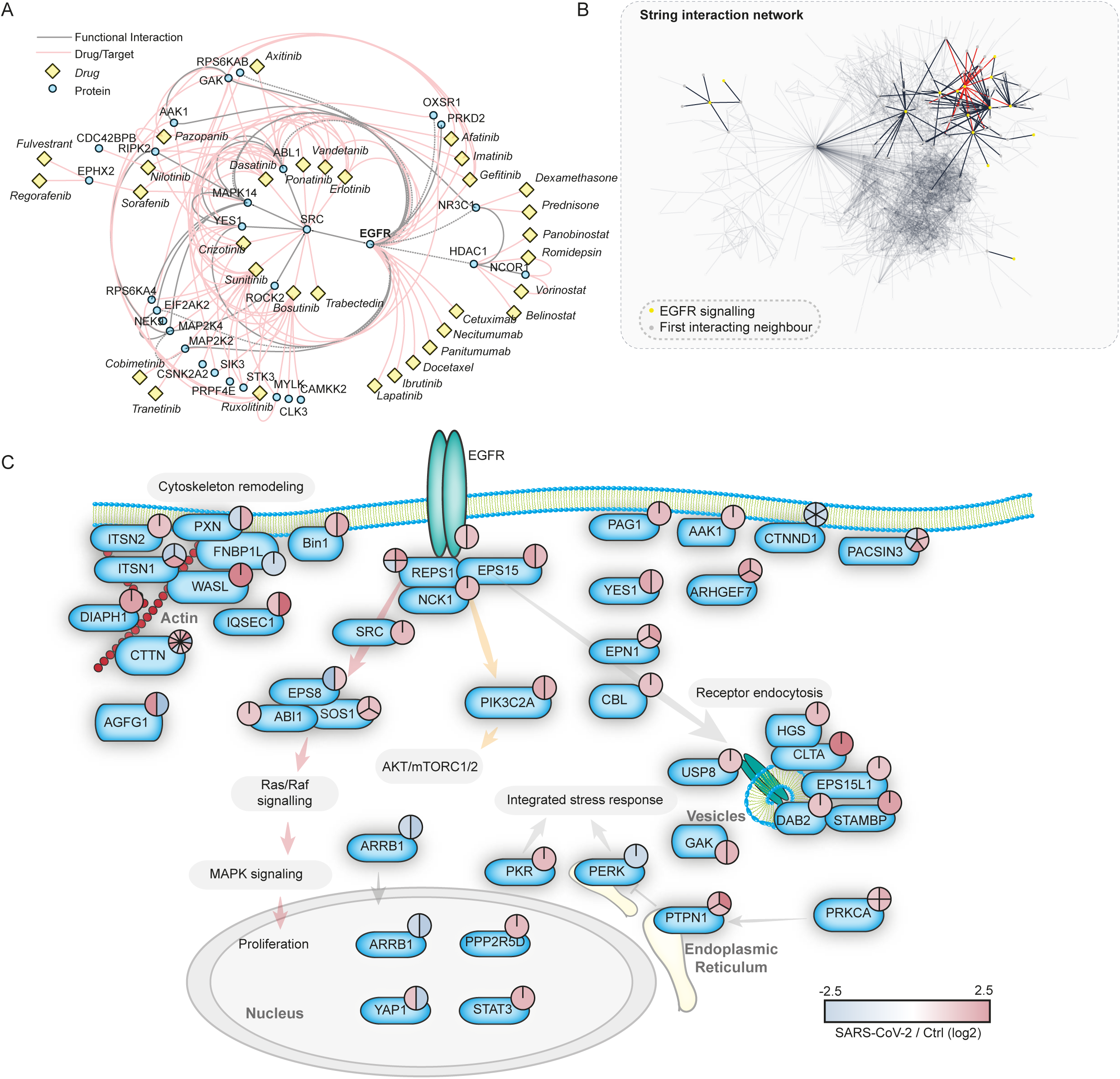
Drug-target phosphoprotein network analysis identifies growth factor signaling as central hub for possible intervention by repurposed drugs. (**A**) Proteins significantly increased in phosphorylation (FC > 1, FDR < 0.05) were subjected to ReactomeFI pathway analysis and overlaid with a network of FDA-approved drugs. The network was filtered for drugs and drug targets only, to identify pathways that could be modulated by drug repurposing. Red lines indicate drug-target interactions, grey lines protein-protein interactions. Identified drugs are represented with yellow rectangles, while proteins are represented by blue circles. (**B**) Search across all proteins with significant phosphorylation changes upon SARS-CoV-2 infection for proteins related to the EGFR pathway. STRING network highlighting all proteins annotated for EGFR signaling and their direct interaction neighbors. Red lines indicate direct EGFR interactions, black lines indicate interactions between pathway members. Grey lines represent filtered interactions to represent the whole network. (**C**) Pathway representation of proteins identified in (B) to be direct functional interactors of EGFR, according to the STRING interaction database (confidence cutoff 0.9). Phosphorylation changes of all significantly regulated sites are indicated by color-coded pie charts. Red indicates upregulation and blue indicates down-regulation.

### Inhibition of growth factor signaling prevents viral replication

Since GFR signaling seemed to be central during SARS-CoV-2 infection, we examined the use of inhibitors as antiviral agents. Since there are several GFRs integrating their signaling and regulating a number of processes inside the cell, directly targeting downstream signaling components is likely to be more successful to prevent signaling of different GFRs and to avoid mixed effects of multiple pathways. GFR signaling, amongst others, results in activation of 1) the RAF/MEK/ERK MAPK signaling cascade and 2) integrates (via phosphoinositide 3 kinase [PI3K] and protein kinase B [AKT]) into mTORC1 signaling to regulating proliferation (Figure 4A). To explore the antiviral efficiency of targeting proteins downstream of GFRs, we first tested the PI3K inhibitors pictilisib and omipalisib (Ippolito et al., 2016; Sarker et al., 2015; Schmid et al., 2016). Both compounds inhibited viral replication, based on their propensity to prevent cytopathogenic effect (CPE) and viral RNA production in cells (Figure 4B-D, S5, and S6).

**Fig. 4.**
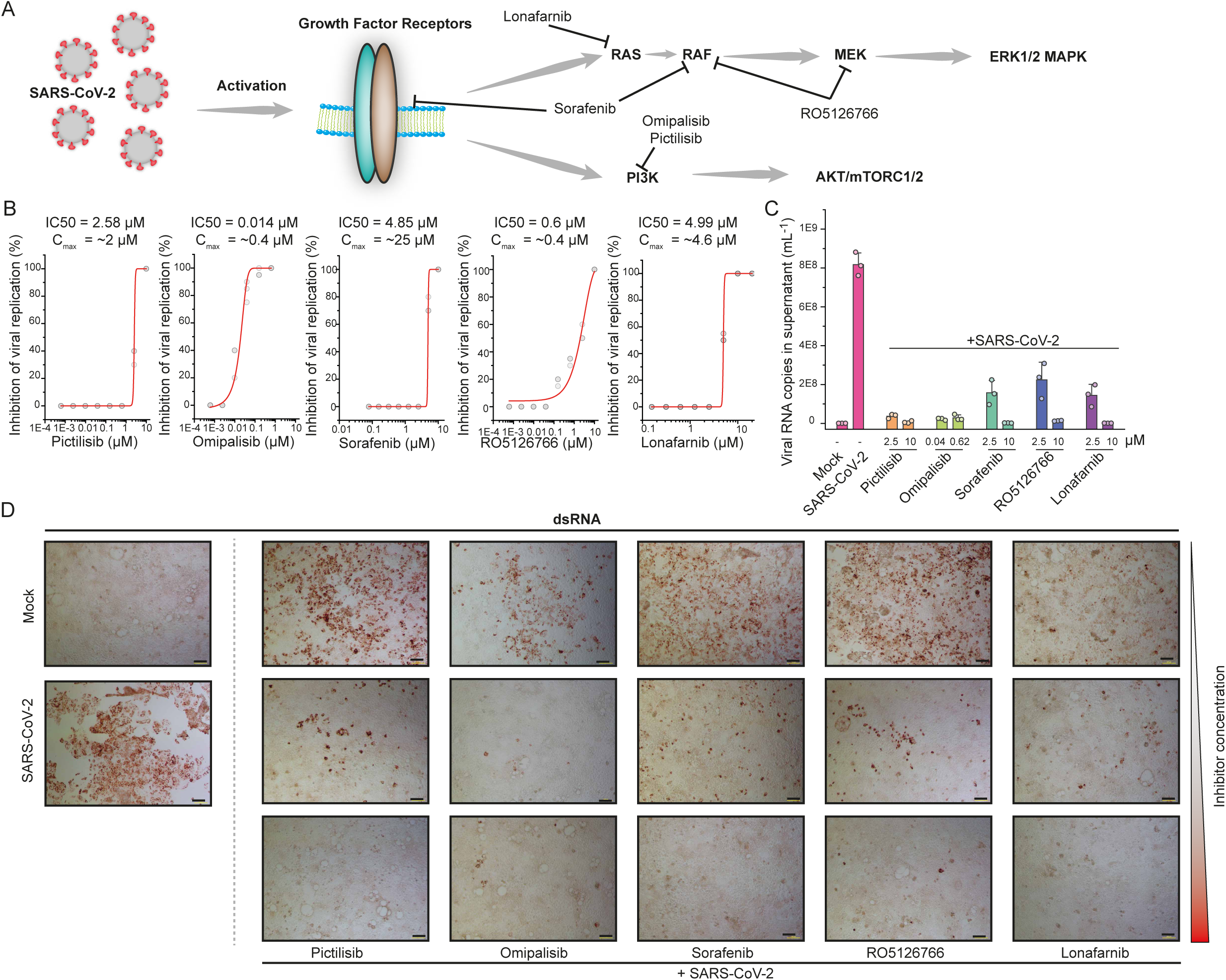
Inhibition of growth factor receptor downstream signaling prevents SARS-CoV-2 replication. (**A**) Schematic representation of growth factor signaling pathways activated upon SARS-CoV-2 infection. Inhibitors tested are indicated and their targets shown. (**B**) Viral replication assay. Percentage inhibition of cytopathic effects (CPE) is plotted versus compound concentration (n = 3 for all compounds). Grey dots indicate replicate measurements, red lines dose-response curve fits. (**C**) Quantification of viral RNA in the supernatant. Supernatant of control cells, infected cells and infected cells treated either with Pictilisib, Omipalisib, Sorafenib, RO5126766 or Lorafenib at indicated concentrations was analyzed by quantitative PCR for viral genome. N = 3, bar indicates mean of replicates, error bars indicate SD. (**D**) Microscopy pictures showing staining for dsRNA to determine viral dsRNA production and CPE. Mock or SARS-CoV-2 infected cells are shown on the left. SARS-CoV-2 infected cells were treated with different concentrations of inhibitors (as indicated) and imaged after 24 hours. Pictilibsib: 0.625 μM, 2.5 μM, 10 μM; omipalisib: 0.01 μM, 0.625 μM, 2.5 μM; sorafenib: 2.5 μM, 5μM, 10 μM; RO5126766: 2.5 μM, 5 μM, 10 μM; lonafarnib: 0.6 μM, 2.5 μM, 10 μM. N = 3, one representative pictures are shown, two more replicates are shown in Figure S4. Scale bar represents 100 μm.

Our drug-target analyses identified mitogen activated protein kinase kinase (MAP2K2, better known as MEK) and the RAF inhibitor sorafenib (Wilhelm et al., 2006) as promising targets inhibiting downstream signaling of GFRs (Figure 4A). Thus, we tested sorafenib and the dual RAF/MEK inhibitor RO5126766 in our viral replication assays. Both compounds inhibited cytopathic effects during infection and the viral replication (Figure 4B, C, D, S5, and S6). Overall, five compounds, inhibiting downstream signaling of GFRs, prevented SARS-CoV-2 replication at clinically achievable concentrations (Figure 4B and 5) (Eskens et al., 2001; Fucile et al., 2014; Martinez-Garcia et al., 2012; Munster et al., 2016; Sarker et al., 2015), emphasizing the importance of GFR signaling during SARS-CoV-2 infection and revealing clinically available treatment options as drug candidates for COVID-19.

**Fig. 5.**
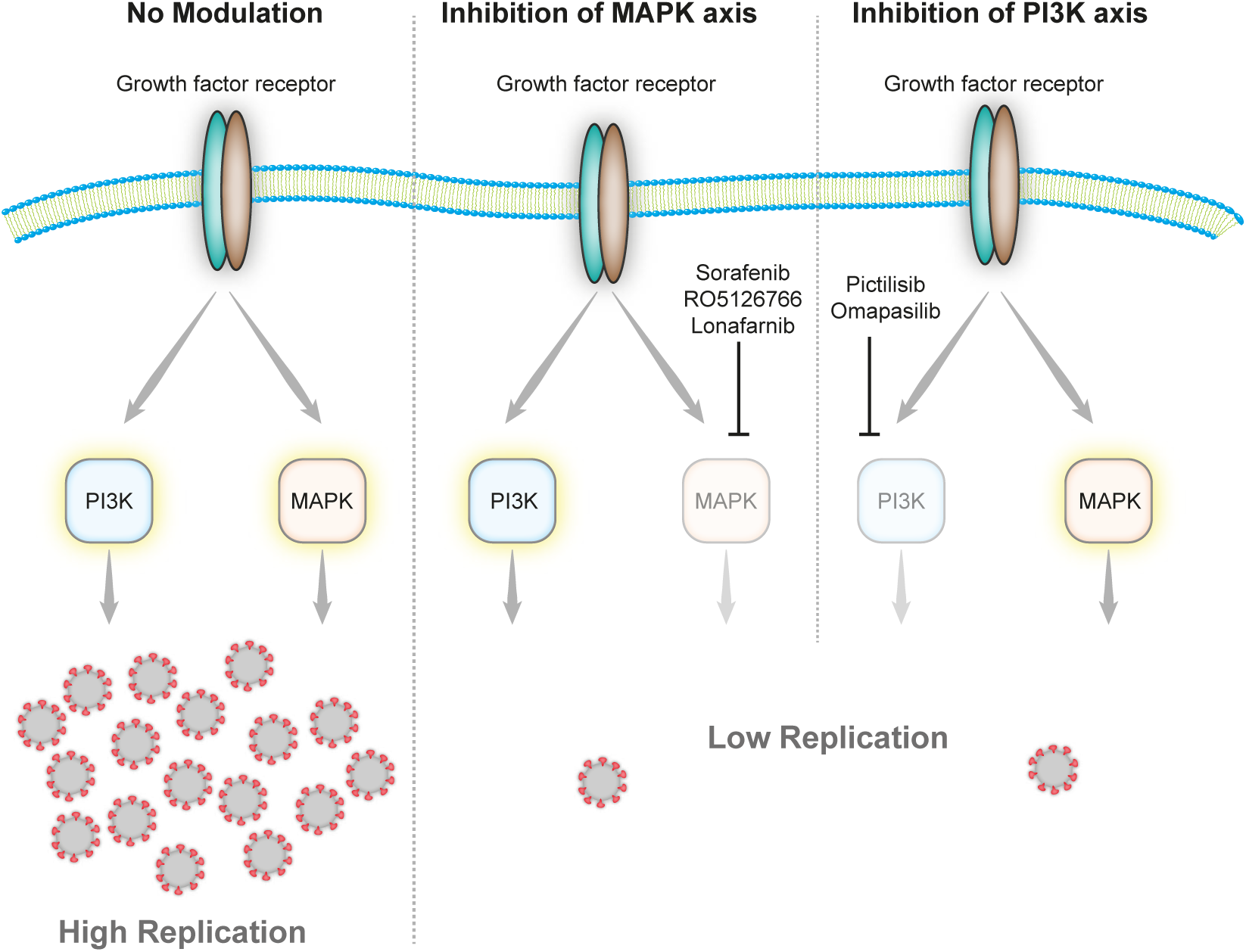
Effect of growth factor signaling on SARS-CoV-2 replication. Upon infection growth factor signaling is activated and leads among others to the induction of Phosphoinositol 3 Kinase (PI3K) and Mitogen Activated Protein Kinase (MAPK) signaling events. Inhibition of either axis of the two (by Sorafenib, RP5126766, Lonafarnib, Pictilisib or Omapalisib) leads to decreased replication of SARS-CoV-2 inside the host cell.

## Discussion

With the rapid spreading of the COVID-19 pandemic, investigating the molecular mechanisms underlying SARS-CoV-2 infection are of high importance. In particular, the processes underlying infection and host-cell response remain unclear. These would offer potential avenues for pharmacological treatment of COVID-19. Here, we report global, differential phosphorylation analysis of host cells after infection with intact SARS-CoV-2 virus. We could identify phosphorylation sites on numerous viral proteins in cells, showing that they can undergo efficient modification in infected cells. Until now, we can only speculate about the host kinases involved and the functions driven by PTMs, which will be an important topic for follow-up studies. For SARS-CoV-1, it was shown that modification of viral proteins can lead to regulation of RNA binding of the nucleoprotein (Wu et al., 2009) and is needed for viral replication. Although similar effects in SARS-CoV-2 are likely, this remains to be studied in this novel virus. A recent paper analyzed the interaction profile of SARS-CoV-2 proteins expressed in HEK293T cells (Gordon et al., 2020). For the heavily phosphorylated nucleoprotein they could identify interactions the host casein kinases, which might indicate possible modification events by the latter. Also for the ORF9b/protein 9b that we found modified in cells, interaction mapping identified MARK kinases as interaction partners.

By exploring the signaling changes inside the host cell, we could gain important insights into host cell signaling during infection. We found essential GFR signaling pathways activated such as EGFR or PDGFR, together with a plethora of RhoGTPase associated signaling molecules. We could furthermore show modulation of the splicing machinery, in line with previous results indicating dependency of viral *in vitro* pathology on the host spliceosome (Bojkova et al., 2020). The same is true for metabolic reprogramming, where we could find differential post-translational modification of most members of the carbon metabolic pathways, namely glycolysis, pentose phosphate and TCA cycle.

A number of drugs to treat COVID-19 have been suggested, largely based on bioinformatics analyses of genetics or cellular data (Gordon et al., 2020; Li et al., 2020; Wang, 2020). However, for many of these compounds, studies explaining their working mechanisms in the context of SARS-CoV-2 or viral assays to determine their efficacy of blocking viral replication in cell models of SARS-CoV-2 infection, are missing. While monitoring signaling changes in host cells, we observed activation of GFR signaling cascades after infection, consistent with other viruses relying on the receptors themselves or elicited signal transduction (Eierhoff et al., 2010; Kung et al., 2011; Lupberger et al., 2011; Ueki et al., 2013; Wu et al., 2017; Zhu et al., 2009). From our data we could not clearly conclude which GFR might be activated and thus tested whether GFR downstream signaling inhibition can prevent SARS-CoV-2 replication, as reported for some other viruses (Baturcam et al., 2019; Pleschka et al., 2001).Previously, temporal kinome analysis identified antiviral potential of RAS/RAF/MEK and PI3K/AKT/ for MERS-CoV (Kindrachuk et al., 2015). By targeting the RAS/RAF/MEK and PI3K/AKT/mTOR downstream axes of GFR signaling, we found efficient inhibition of cytopathic effects and cell destruction (Figure 4). GFR signaling has already been shown to play a role in diverse virus infections as well as in fibrosis induction by SARS-CoV-1 (Beerli et al., 2019; Kung et al., 2011; Lupberger et al., 2011; Pleschka et al., 2001; Ueki et al., 2013; Venkataraman et al., 2017). Thus, our results in cytopathic effects might indeed indicate cytoprotective roles for GFR signaling axes during SARS-CoV-2 infection and possible development of fibrosis (Luo et al., 2020). Notably, some inhibitors used in our study such as omipalisib were already shown to suppress fibrosis progression in patients with idiopathic pulmonary fibrosis, which may share deregulation of signaling pathways involved in lung fibrosis of coronavirus patients (Venkataraman et al., 2017). These findings suggest that inhibitors of GFR downstream signaling may bring benefit to COVID-19 patients independently of their antiviral activity.

Taken together, this study provides new insights into molecular mechanisms elicited by SARS-CoV-2 infection. Proteomic analyses revealed several pathways that are rearranged during infection and showed that targeting of those pathways is a valid strategy to inhibit cytopathic effects triggered by infection.

## Acknowledgements

We thank the Quantitative Proteomics Unit (IBC2, Goethe University Frankfurt) for support and expertise on LC-MS instrumentation and data analysis and Christiane Pallas and Lena Stegmann for support of with experimental work.

## Funding

C.M. was supported by the European Research Council under the European Union’s Seventh Framework Programme (ERC StG 803565), CRC1177 and the Emmy Noether Programme of the Deutsche Forschungsgemeinschaft (DFG, MU 4216/1-1), the Johanna Quandt Young Academy at Goethe, and Aventis Foundation Bridge Award. J.C. was funded by the Hilfe für krebskranke Kinder Frankfurt e.V. and the Frankfurter Stiftung für krebskranke Kinder.

## Author contributions

Conceptualization, K.K., D.B., C.M., and J.C.; Methodology, K.K. and D.B.; Software, K.K.; Formal Analysis, K.K. and G.T.; Investigation, K.K., D.B., and J.C.; Resources, S.C., C.M., and J.C.; Data Curation, K.K.; Writing – Original Draft, K.K. and C.M.; Writing – Review and Editing, K.K., D.B., G.T., C.M., and J.C.; Visualization, K.K.; Supervision, C.M. and J.C.; Project Administration, C.M. and J.C.; Funding Acquisition, S.C., C.M., and J.C.

## Competing interest

The authors filed a patent application on the use of GFR signaling inhibitors for the treatment of COVID-19.

## STAR methods

### Experimental model and subject details

#### Cell culture

Human Caco-2 (Caucasian male) cells, derived from colon carcinoma, was obtained from the Deutsche Sammlung von Mikroorganismen und Zellkulturen (DSMZ; Braunschweig, Germany). Cells were grown at 37°C in Minimal Essential Medium (MEM) supplemented with 10% fetal bovine serum (FBS) and containing 100 IU/ml penicillin and 100 μg/ml streptomycin. All culture reagents were purchased from Sigma.

#### Virus preparation

SARS-CoV-2 was isolated from samples of travelers returning from Wuhan (China) to Frankfurt (Germany) using human colon carcinoma cell line CaCo-2 as described previously^12^. SARS-CoV-2 stocks used in the experiments had undergone one passage on CaCo-2 cells and were stored at –80° C. Virus titers were determined as TCID50/ml in confluent cells in 96-well microtiter plates.

### Method details

#### Antiviral and cell viability assays

Confluent layers of CaCo-2 cells in 96-well plates were infected with SARS-CoV-2 at MOI 0.01. Virus was added together with drugs and incubated in MEM supplemented with 2% FBS with different drug dilutions. Cytopathogenic effect (CPE) was assessed visually 48 h after infection. To assess effects of drugs on Caco-2 cell viability, confluent cell layers were treated with different drug concentration in 96-well plates. The viability was measured using the Rotitest Vital (Roth) according to manufacturer’s instructions. Data for each condition was collected for at least three biological replicates. For dose response curves, data was fitted with all replicates using OriginPro 2020 with the following equation:

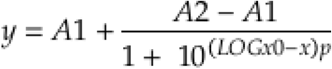

IC50 values were generated by OriginPro 2020 together with metrics for curve fits.

#### Detection of viral replication

Effect of selected compounds on viral replication was assessed by staining of double-stranded RNA, which has been shown to be sufficient for measurement of SARS-CoV-1 replication(Weber et al., 2006). Briefly, cells were fixed with acetone/methanol (40:60) solution 48 h post infection. Immunostaining was performed using a monoclonal antibody directed against dsRNA (1:150 dilution, SCICONS J2, mouse, IgG2a, kappa chain, English & Scientific Consulting Kft., Szirák, Hungary), which was detected with biotin-conjugated secondary antibody (1:1000 dilution, Jackson ImmunoResearch) followed by application streptavidin, peroxidase conjugate (1:3000 dilution, Sigma Aldrich). Lastly, the dsRNA positive cells were visualized by addition of AEC substrate.

#### Sample preparation for mass spectrometry

The sample preparation was performed as described previously(Klann et al., 2020). Briefly, lysates were precipitated by methanol/chloroform and proteins resuspended in 8 M Urea/10 mM EPPS pH 8.2. Concentration of proteins was determined by Bradford assay and 300 μg of protein per samples was used for digestion. For digestion, the samples were diluted to 1 M Urea with 10mM EPPS pH 8.2 and incubated overnight with 1:50 LysC (Wako Chemicals) and 1:100 Sequencing grade trypsin (Promega). Digests were acidified using TFA and tryptic peptideswere purified by tC18 SepPak (50 mg, Waters). 125 μg peptides per sample were TMT labelled and the mixing was normalized after a single injection measurement by LC-MS/MS to equimolar ratios for each channel. 250 μg of pooled peptides were dried for offline High pH Reverse phase fractionation by HPLC (whole cell proteome) and remaining 1 mg of multiplexed peptides were used for phospho-peptide enrichment by High-Select Fe-NTA Phosphopeptide enrichment kit (Thermo Fisher) after manufacturer`s instructions. After enrichment, peptides were dried and resuspended in 70% acetonitrile/0.1% TFA and filtered through a C8 stage tip to remove contaminating Fe-NTA particles. Dried phospho-peptides then were fractionated on C18 (Empore) stage-tip. For fractionation C18 stagetips were washed with 100% acetonitrile twice, followed by equilibration with 0.1% TFA solution. Peptides were loaded in 0.1% TFA solution and washed with water. Elution was performed stepwise with different acetonitrile concentrations in 0.1% Triethylamine solution (5%, 7.5%, 10%, 12.5%, 15%, 17.5%, 20%, 50%). The eight fractions were concatenated into four fractions and dried for LC-MS.

#### Offline high pH reverse phase fractionation

Peptides were fractionated using a Dionex Ultimate 3000 analytical HPLC. 250 μg of pooled and purified TMT-labeled samples were resuspended in 10 mM ammonium-bicarbonate (ABC), 5% ACN, and separated on a 250 mm long C18 column (X-Bridge, 4.6 mm ID, 3.5 μm particle size; Waters) using a multistep gradient from 100% Solvent A (5% ACN, 10 mM ABC in water) to 60% Solvent B (90% ACN, 10 mM ABC in water) over 70 min. Eluting peptides were collected every 45 s into a total of 96 fractions, which were cross-concatenated into 24 fractions and dried for further processing.

#### Liquid chromatography mass spectrometry

All mass spectrometry data was acquired in centroid mode on an Orbitrap Fusion Lumos mass spectrometer hyphenated to an easy-nLC 1200 nano HPLC system using a nanoFlex ion source (ThermoFisher Scientific) applying a spray voltage of 2.6 kV with the transfer tube heated to 300°C and a funnel RF of 30%. Internal mass calibration was enabled (lock mass 445.12003 m/z). Peptides were separated on a self-made, 32 cm long, 75μm ID fused-silica column, packed in house with 1.9 μm C18 particles (ReproSil-Pur, Dr. Maisch) and heated to 50°C using an integrated column oven (Sonation). HPLC solvents consisted of 0.1% Formic acid in water (Buffer A) and 0.1% Formic acid, 80% acetonitrile in water (Buffer B).

For total proteome analysis, a synchronous precursor selection (SPS) multi-notch MS3 method was used in order to minimize ratio compression as previously described (McAlister et al., 2014). Individual peptide fractions were eluted by a non-linear gradient from 7 to 40% B over 90 minutes followed by a step-wise increase to 95% B in 6 minutes which was held for another 9 minutes. Full scan MS spectra (350-1400 m/z) were acquired with a resolution of 120,000 at m/z 200, maximum injection time of 100 ms and AGC target value of 4 × 10^5^. The 20 most intense precursors with a charge state between 2 and 6 per full scan were selected for fragmentation (“Top 20”) and isolated with a quadrupole isolation window of 0.7 Th. MS2 scans were performed in the Ion trap (Turbo) using a maximum injection time of 50ms, AGC target value of 1.5 × 10^4^ and fragmented using CID with a normalized collision energy (NCE) of 35%. SPS-MS3 scans for quantification were performed on the 10 most intense MS2 fragment ions with an isolation window of 0.7 Th (MS) and 2 m/z (MS2). Ions were fragmented using HCD with an NCE of 65% and analyzed in the Orbitrap with a resolution of 50,000 at m/z 200, scan range of 110-500 m/z, AGC target value of 1.5 ×10^5^ and a maximum injection time of 120ms. Repeated sequencing of already acquired precursors was limited by setting a dynamic exclusion of 45 seconds and 7 ppm and advanced peak determination was deactivated.

For phosphopeptide analysis, each peptide fraction was eluted by a linear gradient from 5 to 32% B over 120 minutes followed by a step-wise increase to 95% B in 8 minutes which was held for another 7 minutes. Full scan MS spectra (350-1400 m/z) were acquired with a resolution of 120,000 at m/z 200, maximum injection time of 100 ms and AGC target value of 4 × 10^5^. The 20 most intense precursors per full scan with a charge state between 2 and 5 were selected for fragmentation (“Top 20”), isolated with a quadrupole isolation window of 0.7 Th and fragmented via HCD applying an NCE of 38%. MS2 scans were performed in the Orbitrap using a resolution of 50,000 at m/z 200, maximum injection time of 86ms and AGC target value of 1 × 10^5^. Repeated sequencing of already acquired precursors was limited by setting a dynamic exclusion of 60 seconds and 7 ppm and advanced peak determination was deactivated.

#### Mass spectrometry data analysis

Raw files were analyzed using Proteome Discoverer (PD) 2.4 software (ThermoFisher Scientific). Spectra were selected using default settings and database searches performed using SequestHT node in PD. Database searches were performed against trypsin digested Homo Sapiens SwissProt database, SARS-CoV-2 database (Uniprot pre-release). Static modifications were set as TMT6 at the N-terminus and lysines and carbamidomethyl at cysteine residues. Search was performed using Sequest HT taking the following dynamic modifications into account: Oxidation (M), Phospho (S,T,Y), Met-loss (Protein N-terminus), Acetyl (Protein N-terminus) and Met-loss acetyl (Protein N-terminus). For whole cell proteomics, the same settings were used except phosphorylation was not allowed as dynamic modification. For phospho-proteomics all peptide groups were normalized by summed intensity normalization and then analyzed on peptide level. For whole cell proteomics normalized PSMs were summed for each accession and data exported for further use.

### Quantification and statistical analysis

#### Significance testing

Unless otherwise stated significance was tested by unpaired, two-sided students t-tests with equal variance assumed. Resulting P values were corrected using the Benjamini-Hochberg FDR procedure. Adjusted P values smaller/equal 0.05 were considered significant. For phospho-proteomics an additional fold change cutoff was applied (log2 > |1|), while for total protein levels, due to different dynamic range, a fold change cutoff of log 2 > |0.5| was applied.

#### Prediction of kinase motifs

Kinase motifs of phosphopeptides from SARS-CoV-2 proteins were predicted using NetPhos 3.1 (Blom et al., 1999) and GPS 5.0 (stand-alone version) using the fasta-file of the Uniprot pre-release which was also used for the proteomics data analysis.(Blom et al., 2004; Wang et al., 2020). For NetPhos, only Kinases with a score above 0.5 were considered as positive hits. For GPS 5.0, sequences were submitted separately for S/T- and Y-kinases and the score threshold was set to “high”. For the final list in Supplementary Table 3, only the top hits with the highest scores were considered.

#### Protein co-regulation analysis

Z-scores were calculated for each phospho-site and the total protein levels individually. Phosphosites were collapsed by average. For merging phosphorylation and total protein levels Z-scores for collapsed phosphorylation and protein level were added for each condition and replicate. Thus, both negative Z-scores (downregulation) will produce a lower combined Z-score and vice versa two positive Z-scores will produce a larger combined Z-score. Next, Euclidean distance correlation for all possible protein-protein pairs were calculated, taking all conditions and replicates individually into account. Heatmap was then build by Euclidean distance hierarchical clustering of correlation matrix.

#### Pathway enrichment analysis

Pathway enrichment analysis was performed by ReactomeFI cytoscape plugin or by STRING functional enrichment analysis. Both analysis used Reactome database for pathway annotations.

#### Drug-target network analysis

All proteins were loaded into ReactomeFI cytoscape plugin to visualize protein-protein functional interaction network. Next, drugs were overlaid by ReactomeFI and network was filtered for the drugs and the first interacting partners. Layout was calculated by yFilesLayout algorithm.

#### Interaction network analysis

All proteins showing significant regulation were loaded by OmicsVisualizer cytoscape plugin and STRING interaction network was retrieved with a confidence cutoff of 0.9. For EGFR subnetwork, EGFR was selected with first interacting neighbors.

## Data and Code Availability

The mass spectrometry proteomics data have been deposited to the ProteomeXchange Consortium via the PRIDE (Perez-Riverol et al., 2019) partner repository with the dataset identifiers PXD018357.

## Supplementary Table Legends

**Supplementary Table 1: Quantification of phospho-peptide data.**

Shown are Protein UniProt accessions together with gene names and modification sites (numbers in brackets indicate localization confidence). Log2 ratios between mock and SARS-CoV-2 infected cells were calculated together with P values. P values were adjusted using Benjamini Hochberg FDR adjustment. Additionally replicate quantification values are given.

**Supplementary Table 2: Quantifications of total protein levels of SARS-CoV-2 infected cells and control cells**.

Uniprot Accessions, Protein description, number of PSMs used for quantification, gene symbol, replicate quantifications, log2 ratio and adjusted P values are shown. P values were adjusted using Benjamini Hochberg FDR adjustment.

**Supplementary Table 3: Viral modification sites**.

Modified amino acid, position in peptide, site probability, peptide sequence, number of modified PSMs, unmodified PSMs, protein accession, protein description, position in protein and modification motifs are given for all identified viral modification sites. Additionally, results of kinase predictions by NetPhos 3.1 and GPS5 are added.

**Supplementary Table 4: Reactome Pathway enrichment analysis for proteins found belonging to Cluster I**(Figure 3A/B/C).

Reactome pathway, number of genes found in pathway, enrichment FDR and individual genes in pathway are given.

**Supplementary Table 5: Reactome Pathway enrichment analysis for proteins found belonging to Cluster II**(Figure 3A/E/F).

Reactome pathway, number of genes found in pathway, enrichment P values and individual genes in pathway are given.

**Supplementary Table 6: Reactome Pathway enrichment analysis for proteins found belonging to Cluster III**(Figure 3A).

Reactome pathway, number of genes found in pathway, enrichment FDR and individual genes in pathway are given.

**Supplementary Table 7: Reactome Pathway enrichment analysis for proteins found significantly decreased in total protein levels**(Figure 1C).

Reactome pathway, number of genes found in pathway, enrichment FDR and individual genes in pathway are given.

---

**Supplementary Fig. 1.**
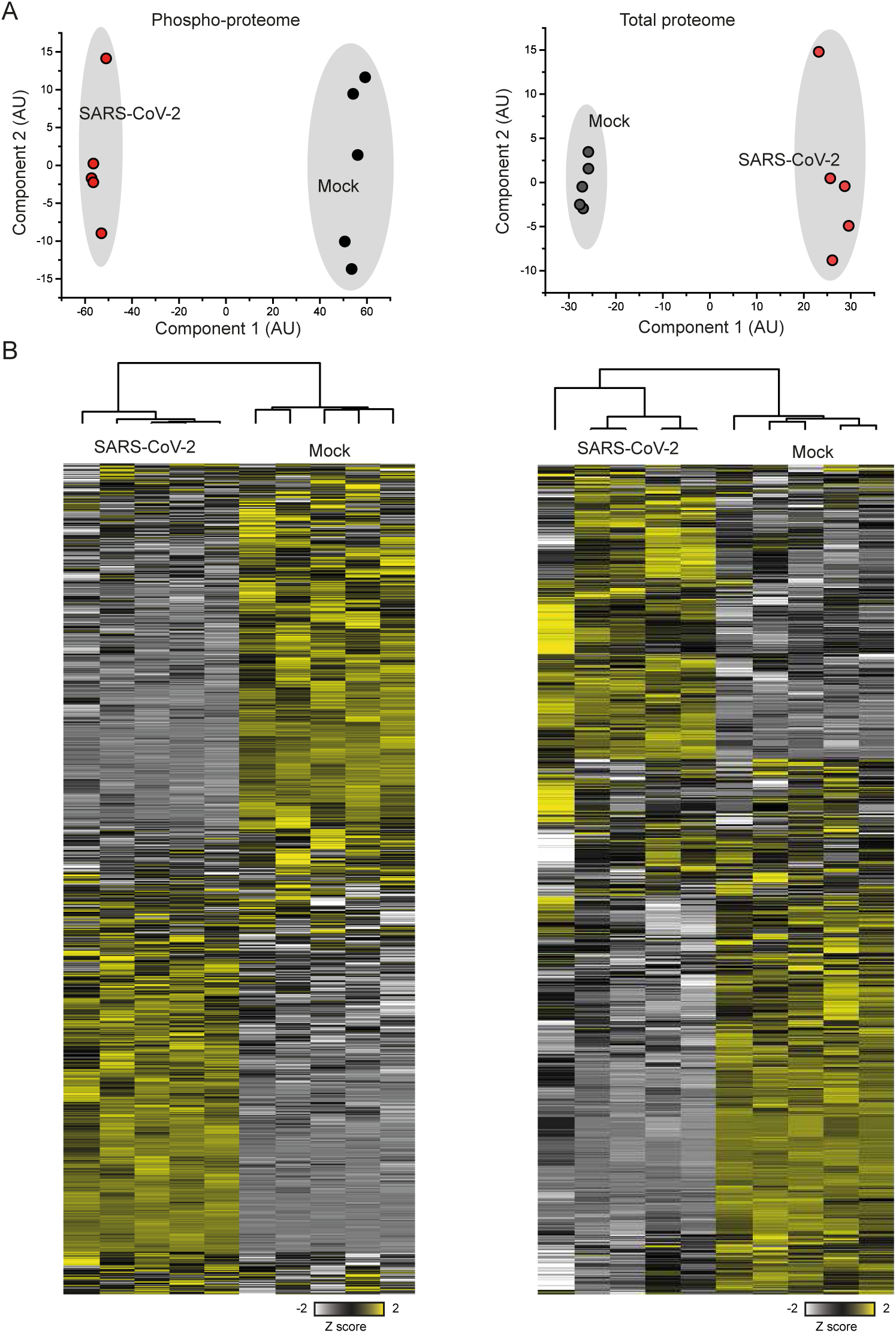
Quality control of proteome datasets. (A) Principal component analyses for phospho- and total proteomes. All quantified phosphopeptides or proteins were log2 transformed and principal component analysis performed in Perseus. Projections were exported and plotted. (B) Heatmaps for phosphoproteome (left) and total proteome (right). All quantified measurements were Z scored and hierarchical clustering carried out with Euclidean distance measure.

**Supplementary Fig. 2.**
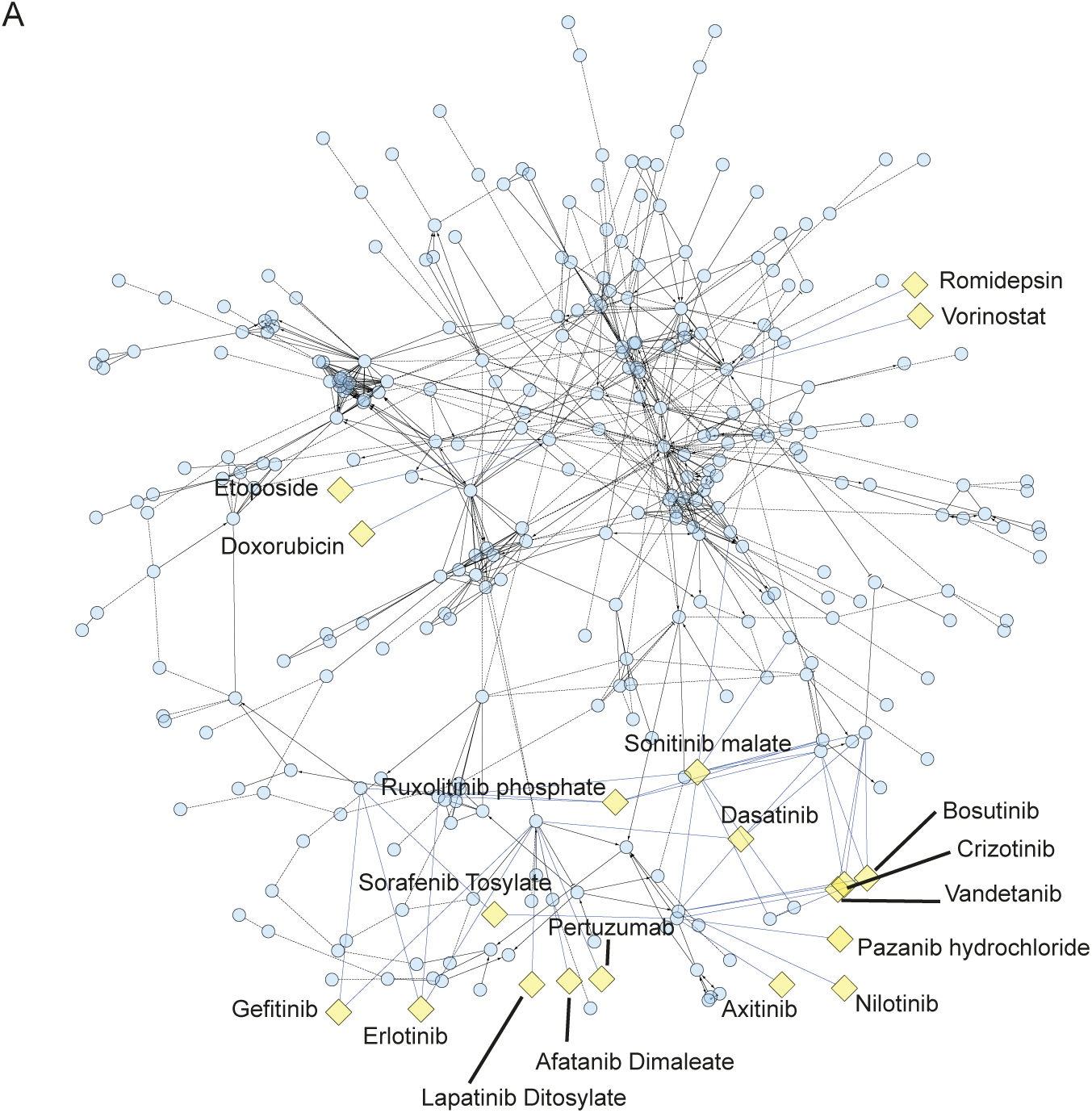
Drug-target network analysis of proteins with significantly decreased phosphorylation. ReactomeFI network was built from all proteins found significantly decreased in phosphorylation (log2 < −1, FDR < 0.05) and overlaid with available drugs. Blue circles indicate proteins, yellow rectangles identified drugs, and lines functional interactions.

**Supplementary Fig. 3:**
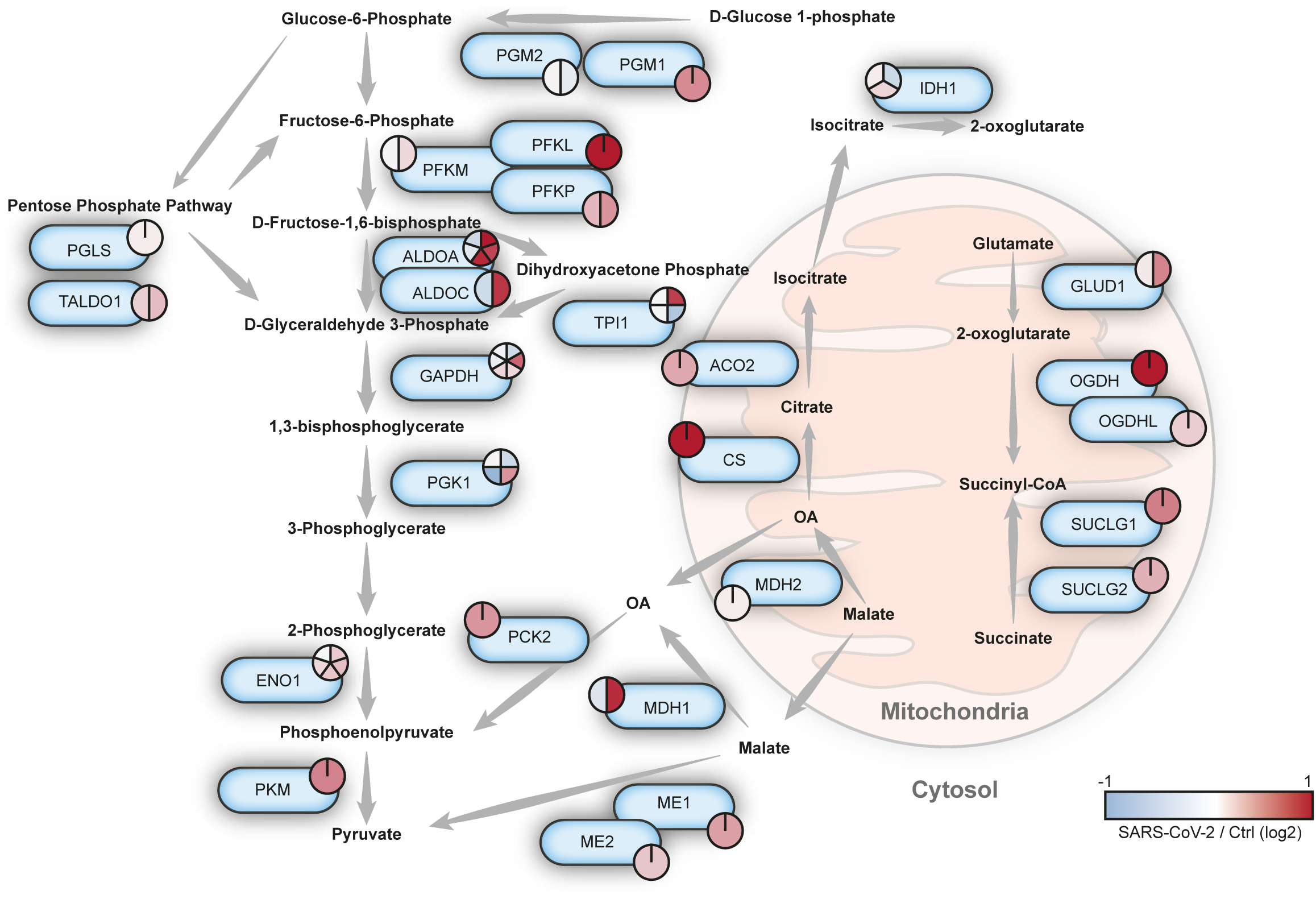
Reprogramming of carbon metabolism upon SARS-CoV-2 infection. Representation of carbon metabolism pathways. All proteins, for which changes in phosphorylation upon SARS-CoV-2 infection could be quantified, were indicated.. Pie charts show fold changes in individual phosphosites, color coded according to the extent to which individual phosphorylation site increased or decreased.

**Supplementary Fig. 4.**
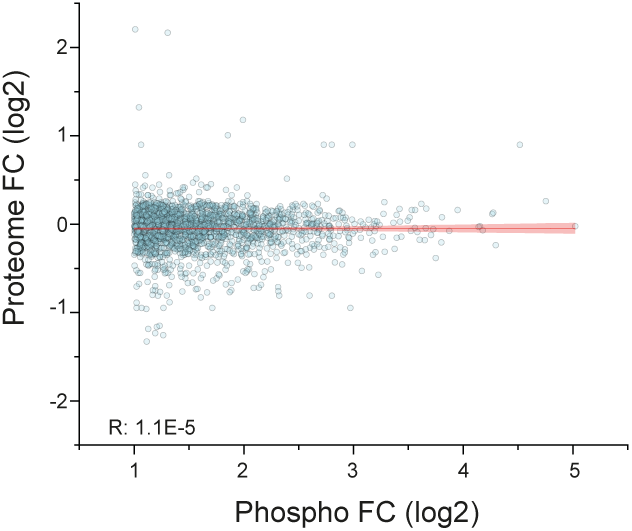
Scatter plot showing phosphopeptide fold changes in comparison to corresponding protein changes for proteins part of the EGFR network. Red line with shade indicates linear fit. No correlation between the two datasets could be observed.

**Supplementary Fig. 5.**
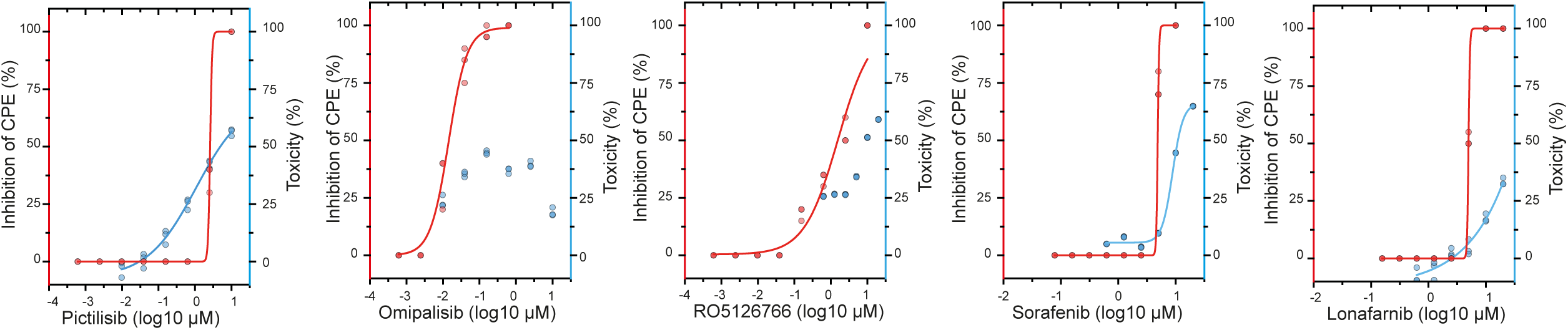
Cytotoxicity data for all tested inhibitors overlaid with CPE data from Figure 4B. Cells were plated and incubated with dose series of different inhibitors. Cytotoxicity was assessed by rotitest vital (n = 3). Red points/axis indicate inhibition of CPE through different inhibitor concentrations. Blue points/axis represent percentage of dead cells compared to control.

**Supplementary Fig. 6.**
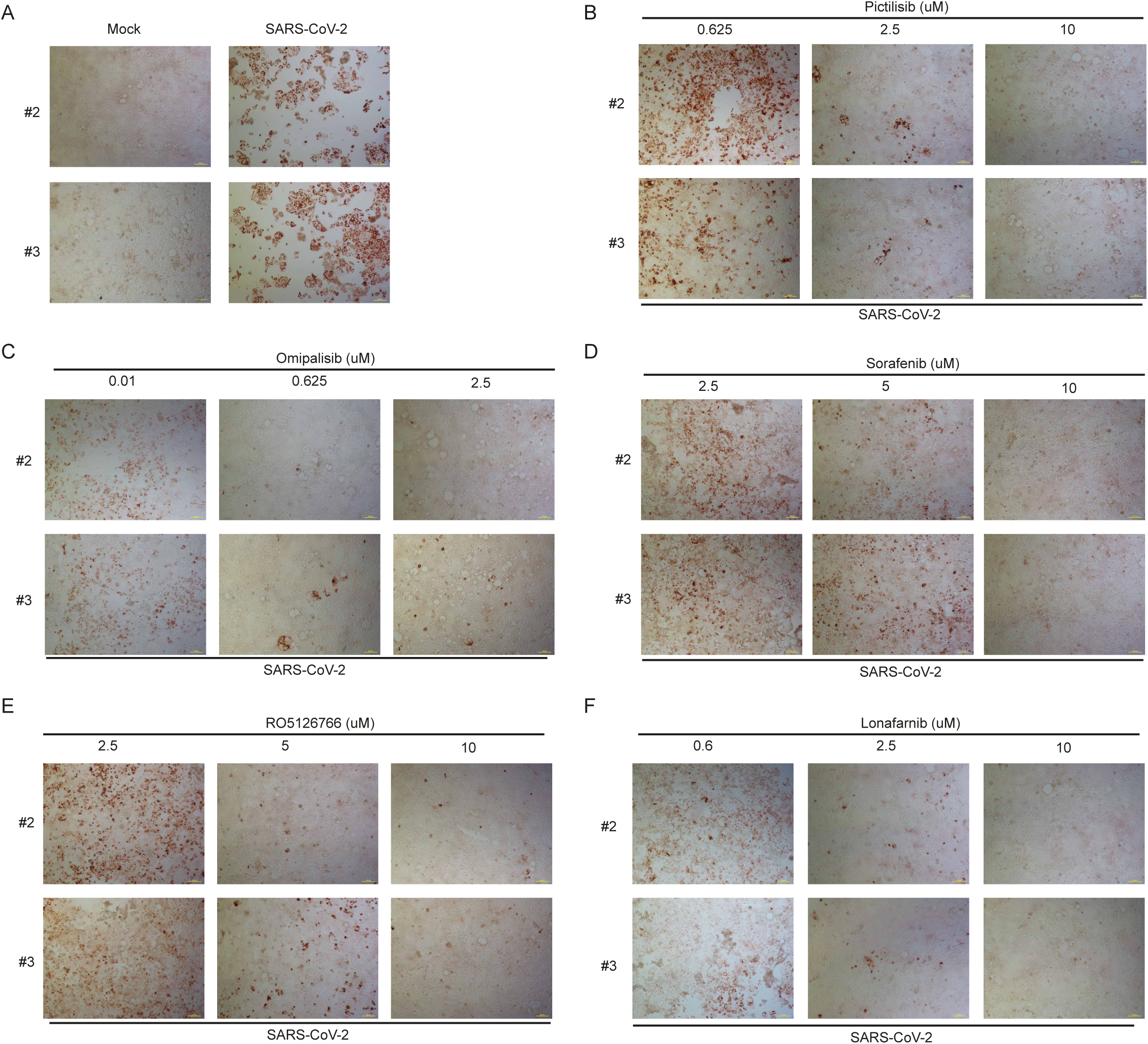
Replicate stainings of dsRNA of SARS-CoV-2 infected cells with and without different inhibitors of GFR signaling. (A) Mock and infected cells after 24 hours of incubation. (B-F) SARS-CoV-2 infected cells with different concentrations of different signaling inhibitors. (B) pictilisib, (C) omipalisib, (D) sorafenib, (E) RO5126766 and (F) lonafarnib. Stainings were performed for dsRNA. Scale bar represents 100 μM.

## Notes

### Summary of Updates

Declaration of interest updated

## References

Baturcam, E., Vollmer, S., Schlüter, H., Maciewicz, R.A., Kurian, N., Vaarala, O., Ludwig, S., and Cunoosamy, D.M. (2019). MEK inhibition drives anti-viral defence in RV but not RSV challenged human airway epithelial cells through AKT/p70S6K/4E-BP1 signalling. Cell Commun. Signal. 17, 78.

Beerli, C., Yakimovich, A., Kilcher, S., Reynoso, G. V, Fläschner, G., Müller, D.J., Hickman, H.D., and Mercer, J. (2019). Vaccinia virus hijacks EGFR signalling to enhance virus spread through rapid and directed infected cell motility. Nat. Microbiol. 4, 216–225.

Blom, N., Gammeltoft, S., and Brunak, S. (1999). Sequence and structure-based prediction of eukaryotic protein phosphorylation sites1 1Edited by F. E. Cohen. J. Mol. Biol. 294, 1351–1362.

Blom, N., Sicheritz-Pontén, T., Gupta, R., Gammeltoft, S., and Brunak, S. (2004). Prediction of post-translational glycosylation and phosphorylation of proteins from the amino acid sequence. Proteomics 4, 1633–1649.

Bojkova, D., Klann, K., Koch, B., Widera, M., Krause, D., Ciesek, S., Cinatl, J., and Münch, C. (2020). SARS-CoV-2 infected host cell proteomics reveal potential therapy targets. Preprint.

Chen, N., Zhou, M., Dong, X., Qu, J., Gong, F., Han, Y., Qiu, Y., Wang, J., Liu, Y., and Wei, Y. (2020). Epidemiological and clinical characteristics of 99 cases of 2019 novel coronavirus pneumonia in Wuhan, China: a descriptive study. Lancet.

Eierhoff, T., Hrincius, E.R., Rescher, U., Ludwig, S., and Ehrhardt, C. (2010). The epidermal growth factor receptor (EGFR) promotes uptake of influenza A viruses (IAV) into host cells. PLoS Pathog. 6, e1001099–e1001099.

Eskens, F.A.L.M., Awada, A., Cutler, D.L., de Jonge, M.J.A., Luyten, G.P.M., Faber, M.N., Statkevich, P., Sparreboom, A., Verweij, J., Hanauske, A.-R., et al. (2001). Phase I and Pharmacokinetic Study of the Oral Farnesyl Transferase Inhibitor SCH 66336 Given Twice Daily to Patients With Advanced Solid Tumors. J. Clin. Oncol. 19, 1167–1175.

Fucile, C., Marenco, S., Bazzica, M., Zuccoli, M.L., Lantieri, F., Robbiano, L., Marini, V., Di Gion, P., Pieri, G., Stura, P., et al. (2014). Measurement of sorafenib plasma concentration by high-performance liquid chromatography in patients with advanced hepatocellular carcinoma: is it useful the application in clinical practice? A pilot study. Med. Oncol. 32, 335.

Gordon, D.E., Jang, G.M., Bouhaddou, M., Xu, J., Obernier, K., O’Meara, M.J., Guo, J.Z., Swaney, D.L., Tummino, T.A., Huttenhain, R., et al. (2020). A SARS-CoV-2-Human Protein-Protein Interaction Map Reveals Drug Targets and Potential Drug-Repurposing. BioRxiv 2020.03.22.002386.

Grimmler, M., Bauer, L., Nousiainen, M., Körner, R., Meister, G., and Fischer, U. (2005). Phosphorylation regulates the activity of the SMN complex during assembly of spliceosomal U snRNPs. EMBO Rep. 6, 70–76.

Herzog, P., Drosten, C., and Müller, M.A. (2008). Plaque assay for human coronavirus NL63 using human colon carcinoma cells. Virol. J. 5, 138.

Ilan, L., Osman, F., Namer, L.S., Eliahu, E., Cohen-Chalamish, S., Ben-Asouli, Y., Banai, Y., and Kaempfer, R. (2017). PKR activation and eIF2α phosphorylation mediate human globin mRNA splicing at spliceosome assembly. Cell Res. 27, 688–704.

Ippolito, T., Tang, G., Mavis, C., Gu, J.J., Hernandez-Ilizaliturri, F.J., and Barth, M.J. (2016). Omipalisib (GSK458), a Novel Pan-PI3K/mTOR Inhibitor, Exhibits In Vitro Anti-Lymphoma Activity in Chemotherapy-Sensitive and -Resistant Models of Burkitt Lymphoma. Blood 128, 5376.

Kindrachuk, J., Ork, B., Hart, B.J., Mazur, S., Holbrook, M.R., Frieman, M.B., Traynor, D., Johnson, R.F., Dyall, J., Kuhn, J.H., et al. (2015). Antiviral Potential of ERK/MAPK and PI3K/AKT/mTOR Signaling Modulation for Middle East Respiratory Syndrome Coronavirus Infection as Identified by Temporal Kinome Analysis. Antimicrob. Agents Chemother. 59, 1088LP–1099.

Klann, K., Tascher, G., and Münch, C. (2020). Functional Translatome Proteomics Reveal Converging and Dose-Dependent Regulation by mTORC1 and eIF2α. Mol. Cell 77, 2-925.e4.

Kung, C.-P., Meckes, D.G., and Raab-Traub, N. (2011). Epstein-Barr Virus LMP1 Activates EGFR, STAT3, and ERK through Effects on PKCδ. J. Virol. 85, 4399LP–4408.

Li, X., Yu, J., Zhang, Z., Ren, J., Peluffo, A.E., Zhang, W., Zhao, Y., Yan, K., Cohen, D., and Wang, W. (2020). Network Bioinformatics Analysis Provides Insight into Drug Repurposing for COVID-2019.

Luo, W., Yu, H., Gou, J., Li, X., Sun, Y., Li, J., and Liu, L. (2020). Clinical Pathology of Critical Patient with Novel Coronavirus Pneumonia (COVID-19).

Lupberger, J., Zeisel, M.B., Xiao, F., Thumann, C., Fofana, I., Zona, L., Davis, C., Mee, C.J., Turek, M., Gorke, S., et al. (2011). EGFR and EphA2 are host factors for hepatitis C virus entry and possible targets for antiviral therapy. Nat. Med. 17, 589–595.

Martinez-Garcia, M., Banerji, U., Albanell, J., Bahleda, R., Dolly, S., Kraeber-Bodéré, F., Rojo, F., Routier, E., Guarin, E., Xu, Z.-X., et al. (2012). First-in-Human, Phase I Dose-Escalation Study of the Safety, Pharmacokinetics, and Pharmacodynamics of RO5126766, a First-in-Class Dual MEK/RAF Inhibitor in Patients with Solid Tumors. Clin. Cancer Res. 18, 4806LP–4819.

Mathew, R., Hartmuth, K., Möhlmann, S., Urlaub, H., Ficner, R., and Lührmann, R. (2008). Phosphorylation of human PRP28 by SRPK2 is required for integration of the U4/U6-U5 tri-snRNP into the spliceosome. Nat. Struct. Mol. Biol. 15, 435–443.

McAlister, G.C., Nusinow, D.P., Jedrychowski, M.P., Wühr, M., Huttlin, E.L., Erickson, B.K., Rad, R., Haas, W., and Gygi, S.P. (2014). MultiNotch MS3 enables accurate, sensitive, and multiplexed detection of differential expression across cancer cell line proteomes. Anal. Chem.

Mermoud, J.E., Cohen, P.T., and Lamond, A.I. (1994). Regulation of mammalian spliceosome assembly by a protein phosphorylation mechanism. EMBO J. 13, 5679–5688.

Munster, P., Aggarwal, R., Hong, D., Schellens, J.H.M., van der Noll, R., Specht, J., Witteveen, P.O., Werner, T.L., Dees, E.C., Bergsland, E., et al. (2016). First-in-Human Phase I Study of GSK2126458, an Oral Pan-Class I Phosphatidylinositol-3-Kinase Inhibitor, in Patients with Advanced Solid Tumor Malignancies. Clin. Cancer Res. 22, 1932LP–1939.

Perez-Riverol, Y., Csordas, A., Bai, J., Bernal-Llinares, M., Hewapathirana, S., Kundu, D.J., Inuganti, A., Griss, J., Mayer, G., Eisenacher, M., et al. (2019). The PRIDE database and related tools and resources in 2019: improving support for quantification data. Nucleic Acids Res. 47, D442–D450.

Pleschka, S., Wolff, T., Ehrhardt, C., Hobom, G., Planz, O., Rapp, U.R., and Ludwig, S. (2001). Influenza virus propagation is impaired by inhibition of the Raf/MEK/ERK signalling cascade. Nat. Cell Biol. 3, 301–305.

Ren, X., Glende, J., Al-Falah, M., Vries, V. de, Schwegmann-Wessels, C., Qu, X., Tan, L., Tschernig, T., Deng, H., Naim, H.Y., et al. (2006). Analysis of ACE2 in polarized epithelial cells: surface expression and function as receptor for severe acute respiratory syndrome-associated coronavirus. J. Gen. Virol. 87, 1691–1695.

Sarker, D., Ang, J.E., Baird, R., Kristeleit, R., Shah, K., Moreno, V., Clarke, P.A., Raynaud, F.I., Levy, G., Ware, J.A., et al. (2015). First-in-Human Phase I Study of Pictilisib (GDC-0941), a Potent Pan–Class I Phosphatidylinositol-3-Kinase (PI3K) Inhibitor, in Patients with Advanced Solid Tumors. Clin. Cancer Res. 21, 77LP–86.

Schmid, P., Pinder, S.E., Wheatley, D., Macaskill, J., Zammit, C., Hu, J., Price, R., Bundred, N., Hadad, S., Shia, A., et al. (2016). Phase II Randomized Preoperative Window-of-Opportunity Study of the PI3K Inhibitor Pictilisib Plus Anastrozole Compared With Anastrozole Alone in Patients With Estrogen Receptor-Positive Breast Cancer. J. Clin. Oncol. 34, 1987–1994.

Smith, M., and Smith, J.C. (2020). Repurposing Therapeutics for COVID-19: Supercomputer-Based Docking to the SARS-CoV-2 Viral Spike Protein and Viral Spike Protein-Human ACE2 Interface.

Tangudu, C., Olivares, H., Netland, J., Perlman, S., and Gallagher, T. (2007). Severe acute respiratory syndrome coronavirus protein 6 accelerates murine coronavirus infections. J. Virol. 81, 1220–1229.

Ueki, I.F., Min-Oo, G., Kalinowski, A., Ballon-Landa, E., Lanier, L.L., Nadel, J.A., and Koff, J.L. (2013). Respiratory virus–induced EGFR activation suppresses IRF1-dependent interferon λ and antiviral defense in airway epithelium. J. Exp. Med. 210, 1929–1936.

Venkataraman, T., and Frieman, M.B. (2017). The role of epidermal growth factor receptor (EGFR) signaling in SARS coronavirus-induced pulmonary fibrosis. Antiviral Res. 143, 142–150.

Venkataraman, T., Coleman, C., and Frieman, M. (2017). Overactive EGFR Signaling Leads to Increased Fibrosis After SARS-CoV Infection. J. Virol. JVI.00182-17.

Wang, J. (2020). Fast Identification of Possible Drug Treatment of Coronavirus Disease -19 (COVID-19) Through Computational Drug Repurposing Study.

Wang, C., Xu, H., Lin, S., Deng, W., Zhou, J., Zhang, Y., Shi, Y., Peng, D., and Xue, Y. (2020). GPS 5.0: An Update on the Prediction of Kinase-specific Phosphorylation Sites in Proteins. Genomics. Proteomics Bioinformatics.

Weber, F., Wagner, V., Rasmussen, S.B., Hartmann, R., and Paludan, S.R. (2006). Double-stranded RNA is produced by positive-strand RNA viruses and DNA viruses but not in detectable amounts by negative-strand RNA viruses. J. Virol. 80, 5059–5064.

Wilhelm, S., Carter, C., Lynch, M., Lowinger, T., Dumas, J., Smith, R.A., Schwartz, B., Simantov, R., and Kelley, S. (2006). Discovery and development of sorafenib: a multikinase inhibitor for treating cancer. Nat. Rev. Drug Discov. 5, 835–844.

Wu, C.-H., Yeh, S.-H., Tsay, Y.-G., Shieh, Y.-H., Kao, C.-L., Chen, Y.-S., Wang, S.-H., Kuo, T.-J., Chen, D.-S., and Chen, P.-J. (2009). Glycogen synthase kinase-3 regulates the phosphorylation of severe acute respiratory syndrome coronavirus nucleocapsid protein and viral replication. J. Biol. Chem. 284, 2—5239.

Wu, Y., Prager, A., Boos, S., Resch, M., Brizic, I., Mach, M., Wildner, S., Scrivano, L., and Adler, B. (2017). Human cytomegalovirus glycoprotein complex gH/gL/gO uses PDGFR-α as a key for entry. PLoS Pathog. 13.

Yarden, Y. (2001). The EGFR family and its ligands in human cancer: signalling mechanisms and therapeutic opportunities. Eur. J. Cancer 37, 3–8.

Zhu, L., Lee, P., Lee, W., Zhao, Y., Yu, D., and Chen, Y. (2009). Rhinovirus-Induced Major Airway Mucin Production Involves a Novel TLR3-EGFR–Dependent Pathway. Am. J. Respir. Cell Mol. Biol. 40, 610–619.

